# Dissociable signatures of visual salience and behavioral relevance across attentional priority maps in human cortex

**DOI:** 10.1101/196642

**Authors:** Thomas C. Sprague, Sirawaj Itthipuripat, Vy A. Vo, John T. Serences

**Author notes:** **Conflict of interest**: the authors declare no conflict of interest.

## Abstract

Computational models posit that visual attention is guided by activity within spatial maps that index the image-computable salience and the behavioral relevance of objects in the scene. However, the simultaneous influence of these factors on putative neural ‘attentional priority maps’ in human cortex is not well understood. We tested the hypothesis that visual salience and behavioral relevance independently impact the activation profile across retinotopically-organized cortical regions by quantifying attentional priority maps measured in human brains using functional MRI while participants attended one of two differentially-salient stimuli. We find that the topography of activation in priority maps, as reflected in the modulation of region-level patterns of population activity, independently indexed the physical salience and behavioral relevance of each scene element. Moreover, salience strongly impacted activation patterns in early visual areas, whereas later visual areas were dominated by relevance. This suggests that prioritizing spatial locations relies on distributed neural codes containing graded representations of salience and relevance across the visual hierarchy.

**Significance Statement:** Often, it is necessary to orient towards bright, unique, or sudden events in the environment – that is, salient stimuli. However, we can focus processing resources on less salient visual information if it is relevant to the task at hand. We tested a theory which supposes that we represent different scene elements according to both their salience and their relevance in a series of ‘priority maps’ by measuring fMRI activation patterns across the human brain and reconstructing spatial maps of the visual scene under different task conditions. We found that different regions indexed either the salience or the relevance of scene items, but not their interaction, suggesting an evolving representation of salience and relevance across different visual areas.

## Introduction

In a typical visual environment, some portions of the scene are <u>*visually salient*</u> by virtue of their image-computable properties, such as luminance, contrast, color saturation, and local feature contrast (Masciocchi et al., 2009; Parkhurst et al., 2002; Usher and Niebur, 1996). Many computational models emphasize the importance of salience in guiding spatial attention (Itti and Koch, 2001; Koch and Ullman, 1985; Stigchel et al., 2009; Theeuwes, 1994, 2010; Veale et al., 2017; Wolfe, 1994). However, visually salient portions of the scene do not necessarily contain behaviorally relevant information. For example, the flashing lights of a police car are relevant if they appear in the rear-view mirror, but less so if they appear in oncoming traffic. Accordingly, other models emphasize <u>*behavioral relevance*</u> in guiding attention, with relevant objects overriding even highly salient scene elements (Egeth and Yantis, 1997; Folk et al., 1992, 2002; Veale et al., 2017; Yantis and Johnson, 1990).

One unifying framework posits that both visual salience and behavioral relevance dissociably influence the activity profile across retinotopic cortex to compute an <u>*attentional priority*</u> map within each region, which can be read out to guide decisions and actions (**Fig 1**; (Fecteau and Munoz, 2006; Itti and Koch, 2001; Serences and Yantis, 2006). This predicts that the topography of activation across an individual retinotopic map (e.g., V1) will reflect the spatial profile of visual salience and/or behavioral relevance of items throughout the scene, and each region’s priority map will weight these factors to varying degrees. Moreover, because representations of visual salience are more closely aligned with retinal input, we predict that attentional priority maps should transition from salience-driven to relevance-driven across the processing hierarchy.

**Figure 1:**
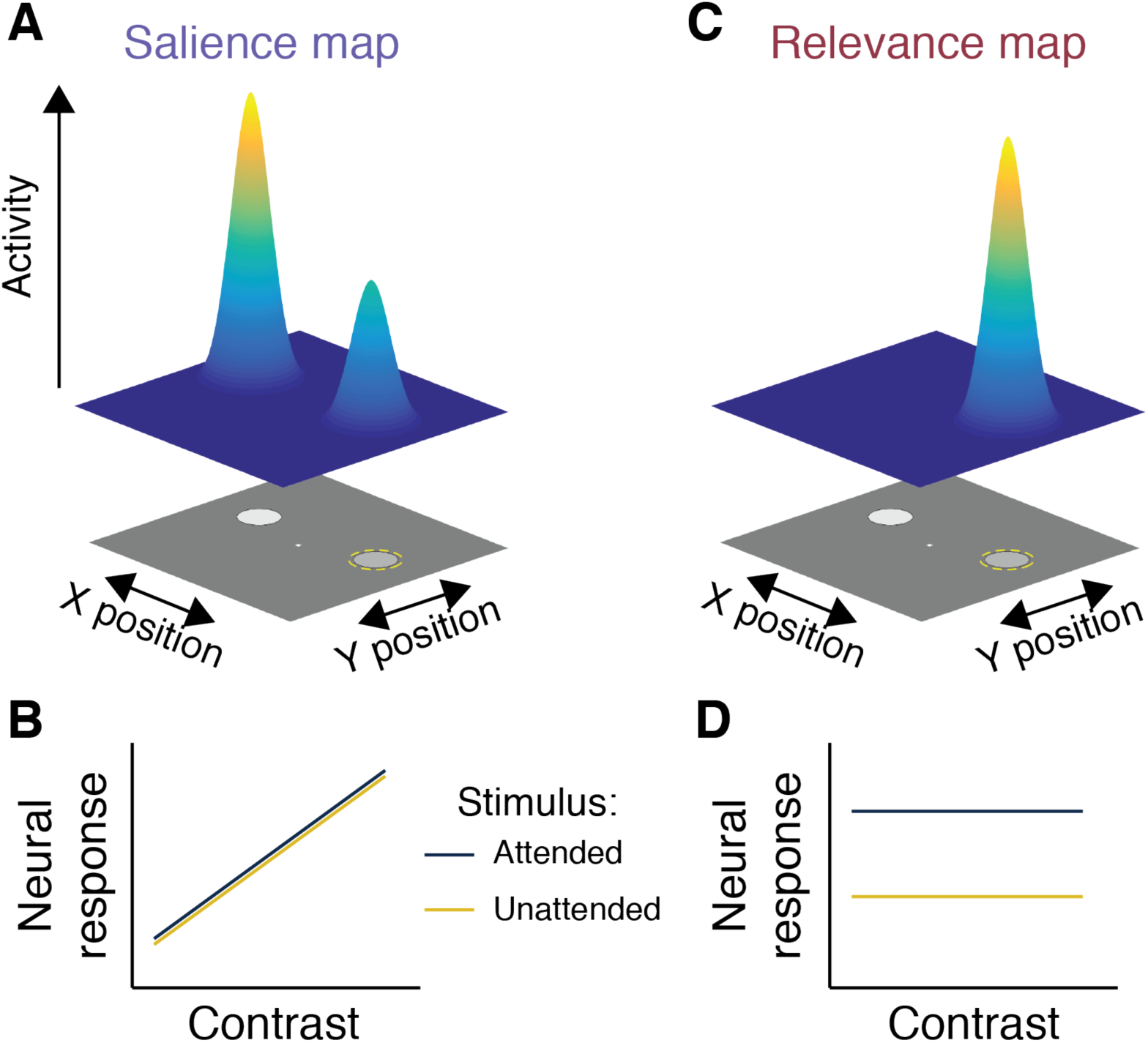
Identifying maps of visual salience and behavioral relevance. **(A)** We define a salience map as a map of the visual scene where each position on the map indexes the importance of the corresponding location within the scene based on image-computable features, such as contrast, motion, or distinctness from the background or other scene items. Accordingly, activity within a salience map should scale with the visual salience of local scene elements. For an example scene, in which two stimuli of differing contrast are presented and a participant is cued to attend to one (dashed yellow circle), a salience map would show higher activation at visual field positions corresponding to higher contrast, even if those elements of the scene are not relevant for behavior. **(B)** For a given location in the scene, activation in a pure salience map would scale only with image-computable features, such as contrast (shown here). **(C)** We define a relevance map as a map of the visual scene where each position on the map indexes the behavioral relevance of the corresponding location. In this example scene, the relevance map would only show activation at locations relevant to the behavior of the observer, independent of their visual salience. This requires that visual locations corresponding to highly-salient but irrelevant stimuli are not reflected in a pure relevance map. **(D)** A location within a pure relevance map would show high activity when the corresponding position is relevant for behavior, and low activity when it is irrelevant. Importantly, a cortical map of visual space can reflect a combination of both visual salience and behavioral relevance by virtue of its position within the visual processing hierarchy.

Previous human neuroimaging studies have identified representations of visual salience in the absence of relevance manipulations (Bogler et al., 2011; Zhang et al., 2012), representations of behavioral relevance under conditions of constant salience (Jerde et al., 2012; Silver et al., 2005; Sprague and Serences, 2013; Vo et al., 2017) and representations of behavioral relevance in the absence of sensory inputs during working memory (Jerde et al., 2012; Rahmati et al.; Saber et al., 2015; Sprague et al., 2014, 2016). While these studies have demonstrated strong preliminary evidence for the existence of representations of attentional priority in retinotopic cortical regions, less is known about how these factors interact when both salience and relevance are parametrically manipulated. For instance, how is an extremely salient, yet behaviorally-irrelevant distractor stimulus represented within region-level attentional priority maps spanning occipital, parietal, and frontal cortex? Previous studies have required subjects to prepare an eye movement or search for a target stimulus (e.g., letter or line) among an array of distractors, with one stimulus marked as salient either by virtue of a unique feature or the abrupt onset of an array element (Balan and Gottlieb, 2006; Bertleff et al., 2016; Gottlieb et al., 1998; Ipata et al., 2006; Thompson et al., 1997). In contrast, here we were interested in manipulating behavioral relevance by cueing one of two stimuli as task-relevant, and manipulating salience by altering the luminance contrast of each stimulus.

Participants attended to one of two objects in a simple visual scene, each with a randomly chosen contrast. We used an inverted encoding model to reconstruct attentional priority maps using each region’s activation pattern to determine how salience and relevance interact to determine representations of attentional priority (**Fig. 1**). We found that early visual areas (e.g., V1) were sensitive to salience, while both earlier and later visual areas (e.g., IPS0) were sensitive to relevance. These results demonstrate a transition from salience-dominated attentional priority maps in early visual cortex to representations dominated by behavioral relevance in higher stages of the visual system.

## Materials & Methods

### Experimental Design

We recruited 8 participants (1 male, all right-handed, 26.5±1.15 yrs age, mean ± SEM), including 1 author (subject ID: AP). Two of these participants had never participated in visual functional neuroimaging experiments before (subject IDs: BA and BF). All others have participated in other experiments in the lab (Ester et al., 2015; Sprague and Serences, 2013; Sprague et al., 2014, 2016; Vo et al., 2017). All procedures were approved by the UCSD Institutional Review Board, all participants gave written informed consent before participating, and all participants were compensated for their time ($20/hr for scanning sessions, $10/hr for behavioral sessions; participant AP, who was an author, was not compensated).

Each participant performed a 1-hr training session before scanning during which they were familiarized with all tasks performed inside the scanner. We also used this session to establish initial behavioral performance thresholds by manipulating task difficulty across behavioral blocks.

We scanned participants for a single 2-hr main task scanning session comprising at least 4 mapping task runs and 4 selective attention task runs (broken into 2 sub-runs each, see below). All participants also underwent additional localizer and retinotopic mapping scanning sessions to independently identify ROIs (see “Region of interest definition”).

We presented stimuli using the Psychophysics toolbox (Brainard, 1997; Pelli, 1997) for MATLAB (The Mathworks, Natick, Mass). During scanning sessions, we rear-projected visual stimuli onto a 110 cm-wide screen placed ~370 cm from the participant’s eyes at the foot of the scanner bore using a contrast-linearized LCD projector (1024×768, 60 Hz). In the behavioral familiarization session, we presented stimuli on a contrast-linearized LCD monitor (1920×1080, 60 Hz) 62 cm from participants, who were comfortably seated in a dimmed room and positioned using a chin rest. For all sessions and tasks (main selective attention task, mapping task, and localizer), we presented all stimuli on a neutral gray 6.82° circular aperture, surrounded by black (only aperture shown in **Fig. 2A**).

**Figure 2:**
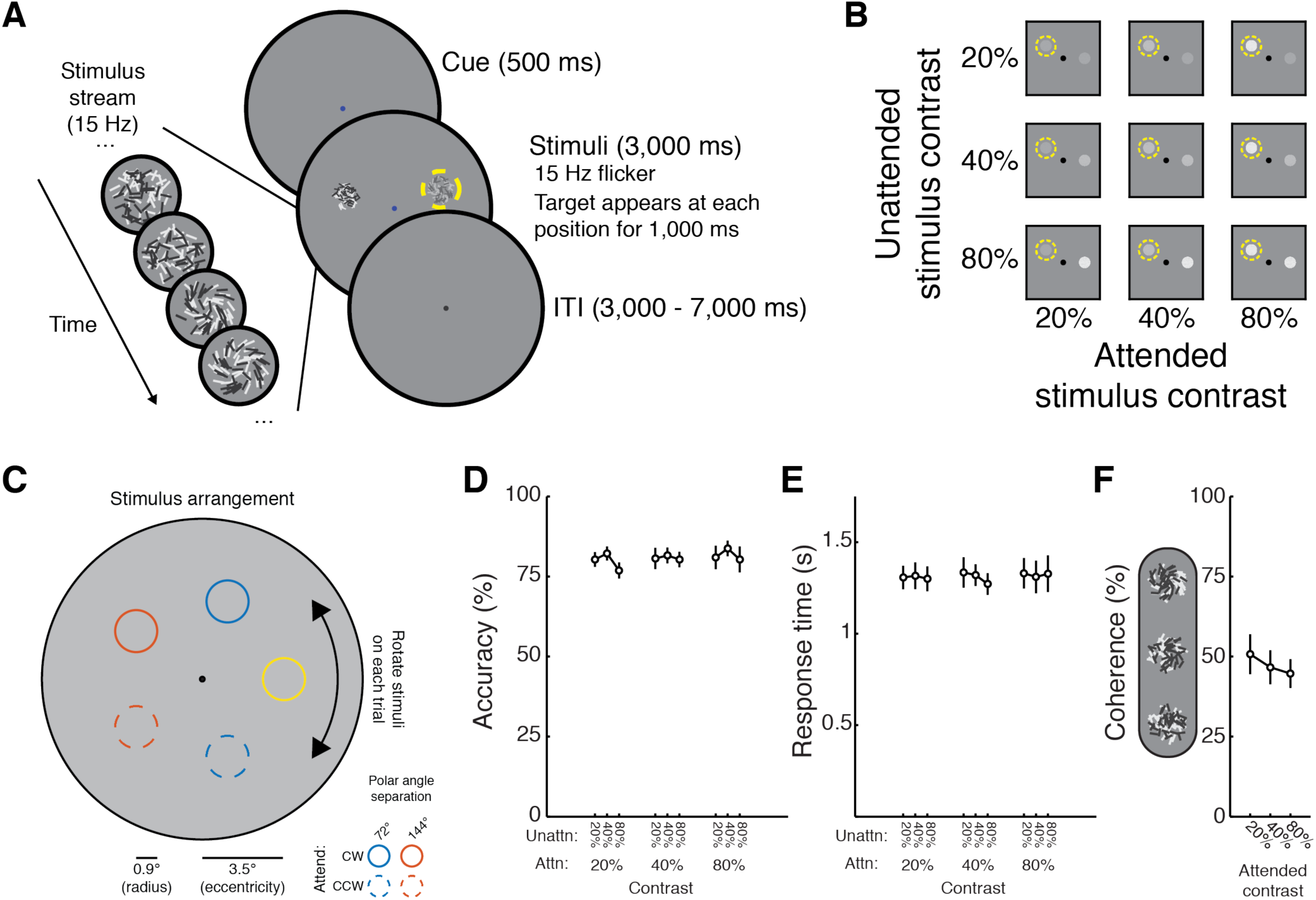
Measuring spatial maps of visual salience and behavioral relevance. **(A)** On each trial, a fixation cue (red or blue dot) indicated whether participants (*n* = 8) should attend the clockwise or counterclockwise of two stimuli which appeared 500 ms after cue onset. We presented stimuli at 3.5° eccentricity separated by 144° or 72° polar angle, and for each trial we randomly rotated the display such that each polar angle was equally-likely to be stimulated on a given trial. Clockwise and anticlockwise are with respect to the polar angle between the two stimuli (this was always unambiguous, as the stimuli were never 180° apart). Stimuli, which consisted of randomly-oriented light and dark lines, flickered at 15 Hz for 3,000 ms (left inset). On every trial, a target appeared within each stimulus stream. Targets (1,000 ms) were variable-coherence spirals oriented clockwise (example shown) or anti-clockwise. Participants responded with a button press to indicate the direction of the spiral in the cued stimulus. **(B)** We presented the attended stimulus and unattended stimulus each at one of three logarithmically-spaced contrast levels, fully crossed. **(C)** On each trial, two stimuli appeared at 3.5° eccentricity, and could be either 72° or 144° polar angle apart. The attended stimulus (yellow for this example arrangement) could be any of the 5 ‘base’ positions. The attended stimulus could appear clockwise (CW; solid lines) or counter-clockwise (CCW; dashed lines) with respect to the unattended stimulus. The entire stimulus display was rotated randomly about fixation on each trial, so each and every trial involved a distinct stimulus display. **(D)** Behavioral performance and **(E)** response time did not reliably vary as a function of attended or unattended stimulus contrast, or their interaction (2-way permuted repeated-measures ANOVA, *p*-values for main effect of attended, unattended contrast and interaction for accuracy: 0.359, 0.096, 0.853; for response time: 0.926, 0.143, 0.705). **(F)** Average coherence used for each contrast to achieve performance in B-C. Though qualitatively the coherence decreases with increasing contrast, there is no main effect of target contrast on mean coherence (1-way permuted repeated-measures ANOVA, *p* = 0.153). Error bars SEM across participants, *n* = 8.

### Selective Attention Task

We instructed participants to attend to one of two random line stimuli (RLS), each appearing at one of three contrasts (20%, 40%, 80%). On each trial, each RLS stimulus would cohere into a ‘spiral’ form and participants responded with one of two button presses indicating which direction of spiral they detected in the cued RLS. We designed this stimulus to minimize the allocation of non-spatial feature-based attention, as well as to minimize the influence of potential radial biases in orientation preference in visual cortex as a function of preferred polar angle (Freeman et al., 2011, 2013).

Each trial began with a 500 ms symbolic attention cue indicating which of the two stimuli to attend, followed by appearance of both stimuli simultaneously. Both stimuli remained onscreen for 3,000 ms, during which time a 1,000 ms spiral target appeared independently at each stimulus position. The target onset was randomly chosen on each trial for each stimulus (attended and unattended) from a uniform distribution spanning 500-1,500 ms.

Both stimuli always appeared along an invisible iso-eccentric ring 3.5° from fixation, and had a radius of 1.05°. On each trial, the two stimuli appeared either 72 or 144 degrees polar angle apart (4.11° and 6.66° distance between centers, respectively). We randomly rotated the stimulus array on each trial (0-72° polar angle) around fixation, so the positions of the stimuli on each trial were entirely unique. The color of the symbolic attention cue presented at fixation indicated with 100% validity whether to attend to the clockwise (blue) or the counterclockwise (red) stimulus (Fig. 2A). Each trial was separated by an inter-trial interval (ITI) drawn from a uniform distribution spanning 2.5-6.5 s at .09 s steps (distribution of ITIs was computed with MATLAB’s linspace command as ITIs=linspace(2.5,6.5,45)), resulting in an average trial duration of 8 s.

Both stimuli flickered in-phase at 15 Hz (2 frames on, 2 frames off, monitor refresh rate of 60Hz). Each stimulus consisted of 35 light and 35 dark lines, each 0.3° long and 0.035° thick, which were replotted during each flicker period centered at random coordinates drawn from a uniform disc 0.9° in radius. On flicker periods with no targets, the orientation of each line was drawn from a uniform distribution. On flicker periods with targets, the orientation of a random subset of lines (defined by the target coherence) was oriented either 45° anti-clockwise or clockwise from radial relative to the stimulus center (see Fig. 2A). Participants responded with a left button press to indicate targets oriented anticlockwise relative to radial, and a right button press indicated those clockwise from radial.

We counterbalanced the attended and unattended stimuli based on contrast (20%, 40%, or 80%; Fig. 2B), stimulus separation distance (72° or 144° polar angle; Fig. 2C), and approximate position of the attended stimulus (1 of 5 “base” positions). Accordingly, for a single repetition of all trial types, we acquired 90 trials, broken up into 2 sub-runs, each lasting 382 s (45 trials, 8 s each, 12 s blank screen at beginning of scan, 10 s blank screen at end of each scan). Participants performed 8 sub-runs, resulting in a full dataset of 360 trials per participant.

To keep performance below ceiling and at an approximately fixed level (at ~80%), we adjusted the coherence of targets (defined as the percentage of lines forming a spiral target on each flicker cycle; Fig. 2F) independently for 20%, 40%, and 80% contrast targets before the start of each full run (i.e. 2 sub-runs).

### Spatial Mapping Task

To estimate a spatial encoding model for each voxel (see below), we presented a flickering checkerboard stimulus at different positions across the screen on each trial. Participants attended these stimuli to identify rare target events (changes in checkerboard contrast on 10 of 47 (21.2%) of trials, evenly split between increments & decrements). During each run, we chose the position of each trial’s checkerboard (0.9° radius, 70% contrast, 6 Hz full-field flicker) from a triangular grid of 37 possible positions and added a random uniform circular jitter (0.5° radius; **Fig 2C**). As in a previous report, we rotated the “base” position of the triangular grid on each scanner run to increase the spatial sampling density (Sprague et al., 2016). Accordingly, every mapping trial was unique. The base triangular grid of stimulus positions separated stimuli by 1.5°, and extended 4.5° from fixation (3 steps). This, combined with random jitter and the radius of the mapping stimulus, resulted in a visual “field of view” (region of the visual field stimulated) of 5.9° from fixation for our spatial encoding model. On trials in which the checkerboard stimulus overlapped the fixation point, we drew a small aperture around fixation (0.8° diameter).

Each trial consisted of a 3,000 ms stimulus presentation period followed by a 2,000-6,000 ms ITI (uniformly sampled). On trials with targets, the checkerboard was dimmed or brightened for 500 ms, beginning at least 500 ms after stimulus onset and ending at least 500 ms before stimulus offset. We instructed participants to only respond if they detected a change in checkerboard contrast and to minimize false alarms. We discarded all target-present trials when estimating spatial encoding models. To ensure participants performed below ceiling we adjusted the task difficulty between each mapping run by changing the percentage contrast change for target trials. Each run consisted of 47 trials (10 of which included targets), a 12 s blank period at the beginning of the run, and a 10 s blank period at the end of the run, totaling 352 s.

### Visual Attention Localizer Task

To identify voxels responsive to the region of the screen subtended by the mapping and selective attention stimuli, all participants performed several runs of a visual localizer task reported previously (Sprague et al., 2016). Participants performed between 3 and 8 runs of this task in total. For one participant, we used data from the same task, acquired for a different experiment (Sprague et al., 2016) at a different scanning resolution (participant AS, 2×2×3 mm voxel size), resampled to the resolution used here. For four participants, including the one scanned with a different protocol, the entire background of the screen (18.2° by 13.65° rectangle) was gray (no circular aperture).

Each trial consisted of a flickering radial checkerboard hemi-annulus presented on the left or right half of fixation subtending 0.8° to 6.0° eccentricity around fixation (6 Hz contrast reversal flicker, 100% contrast, 10.0 s duration). On each trial, participants performed a demanding spatial working memory (WM) task in which they carefully maintained the position of a target stimulus (red dot presented over the stimulus, 500 ms) over a 3,000 ms delay interval, after which a probe stimulus (green dot, 750 ms) appeared near the remembered target position. Participants reported whether the green probe dot appeared to the left of or right of the remembered position; or above or below the remembered position, as prompted by the appearance of a 1.0°-long bar at fixation (horizontal bar: left vs. right; vertical bar: above vs. below; 1.5 s response window). We manipulated the target-probe separation distance between runs to ensure performance was below ceiling.

WM trials could occur beginning 1.0 s after checkerboard onset and ended at latest 2.5 s before checkerboard offset. All trials were separated by a 3 – 5 s ITI (uniformly spaced across trials), and we included 4 null trials in which no checkerboard or WM stimuli appeared (10 s long each). Each run featured 16 total stimulus-present trials, a 14 s blank screen at the beginning of the run and a 10 s blank screen at the end of the run, totaling 304 s.

### Functional MRI Acquisition

We scanned all participants on a 3 T research-dedicated GE MR750 scanner located at the UCSD Keck Center for Functional Magnetic Resonance Imaging with a 32 channel send/receive head coil (Nova Medical, Wilmington, MA). We acquired functional data using a gradient echo planar imaging (EPI) pulse sequence (19.2 × 19.2 cm field of view, 64 × 64 matrix size, 35 3-mm-thick slices with 0-mm gap, axial orientation, TR = 2,000 ms, TE = 30 ms, flip angle = 90°, voxel size 3 mm isotropic).

To anatomically coregister images across sessions, and within each session, we also acquired a high resolution anatomical scan during each scanning session (FSPGR T1-weighted sequence, TR/TE = 11/3.3 ms, TI = 1,100 ms, 172 slices, flip angle = 18°, 1 mm^3^ resolution). For all sessions but one, anatomical scans were acquired with ASSET acceleration. For the remaining session, we used an 8 channel send/receive head coil and no ASSET acceleration to acquire anatomical images with minimal signal inhomogeneity near the coil surface, which enabled improved segmentation of the gray-white matter boundary. We transformed these anatomical images to Talairach space and then reconstructed the gray/white matter surface boundary in BrainVoyager 2.6.1 (BrainInnovations, The Netherlands) which we used for identifying ROIs.

### FMRI Preprocessing

We preprocessed fMRI data as described in our previous reports (Sprague et al., 2014, 2016). We coregistered functional images to a common anatomical scan across sessions (used to identify gray/white matter surface boundary as described above) by first aligning all functional images within a session to that session’s anatomical scan, then aligning that session’s scan to the common anatomical scan. We performed all preprocessing using FSL (Oxford, UK) and BrainVoyager 2.6.1 (BrainInnovations). Preprocessing included unwarping the EPI images using routines provided by FSL, then slice-time correction, three-dimensional motion correction (six-parameter affine transform), temporal high-pass filtering (to remove first-, second-and third-order drift), transformation to Talairach space (resampling to 3×3×3 mm resolution) in BrainVoyager, and finally normalization of signal amplitudes by converting to Z-scores separately for each run using custom MATLAB scripts. We did not perform any spatial smoothing beyond the smoothing introduced by resampling during the co-registration of the functional images, motion correction and transformation to Talairach space. All subsequent analyses were computed using custom code written in MATLAB (release 2015a).

### Region of Interest Definition

Based on our previous work (Sprague et al., 2014, 2016), we identified 10 *a priori* ROIs using independent scanning runs from those used for all analyses reported in the text. For retinotopic ROIs (V1-V3, hV4, V3A, IPS0-IPS3), we utilized a combination of retinotopic mapping techniques. Each participant completed several scans of meridian mapping in which we alternately presented flickering checkerboard “bowties” along the horizontal and vertical meridians. Additionally, each participant completed several runs of an attention-demanding polar angle mapping task in which they detected brief contrast changes of a slowly-rotating checkerboard wedge (described in detail in (Sprague and Serences, 2013)). We used a combination of maps of visual field meridians and polar angle preference for each voxel to identify retinotopic ROIs (Engel et al., 1994; Swisher et al., 2007). Polar angle maps computed using the attention-demanding mapping task for several participants are available in previous publications (AI: (Sprague and Serences, 2013); AL and AP: (Ester et al., 2015)). We combined left- and right-hemispheres for all ROIs, as well as dorsal and ventral aspects of V2 and V3 for all analyses by concatenating voxels.

We defined superior precentral sulcus (sPCS), the putative human homolog of macaque frontal eye field (FEF; Mackey et al., 2017; Srimal and Curtis, 2008) by plotting voxels active during either the left or right conditions of the localizer task described above (FDR corrected, *q* = 0.05) on the reconstructed gray/white matter boundary of each participant’s brain and manually identifying clusters appearing near the superior portion of the precentral sulcus, following previous reports (Srimal and Curtis, 2008). Though retinotopic maps have previously been reported in this region (Hagler and Sereno, 2006; Mackey et al., 2017), we did not observe topographic visual field representations. We anticipate this is due to the substantially limited visual field of view we could achieve inside the scanner (maximum eccentricity: ~7°).

### Inverted Encoding Model

To reconstruct images of salience and/or relevance maps carried by activation patterns measured over entire regions of interest, we implemented an inverted encoding model (IEM) for spatial position (Sprague and Serences, 2013). This analysis involves first estimating an encoding model (sensitivity profile over the relevant feature dimension(s) as parameterized by a small number of modeled information channels) for each voxel in a region using a “training set” of data reserved for this purpose (four spatial mapping runs). Then, the encoding models across all voxels within a region are inverted to estimate a mapping used to transform novel activation patterns from a “test set” (selective attention task runs) into activation in the modeled set of information channels.

Adopting analysis procedures from previous work, we built an encoding model for spatial position based on a linear combination of spatial filters (Sprague and Serences, 2013; Sprague et al., 2014, 2015). Each voxel’s response was modeled as a weighted sum of 37 identically-shaped spatial filters arrayed in a triangular grid (**Fig. 3**). Centers were spaced by 1.59° and each filter was a Gaussian-like function with full-width half-maximum of 1.75°:

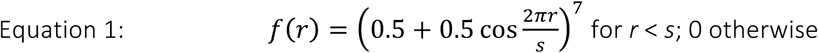

Where *r* is the distance from the filter center and *s* is a “size constant” reflecting the distance from the center of each spatial filter at which the filter returns to 0. Values greater than this are set to 0, resulting in a single smooth round filter at each position along the triangular grid (*s* = 4.404°; see **Fig. 3** for illustration of filter layout and shape; see also (Sprague and Serences, 2013; Sprague et al., 2014, 2016)).

**Figure 3:**
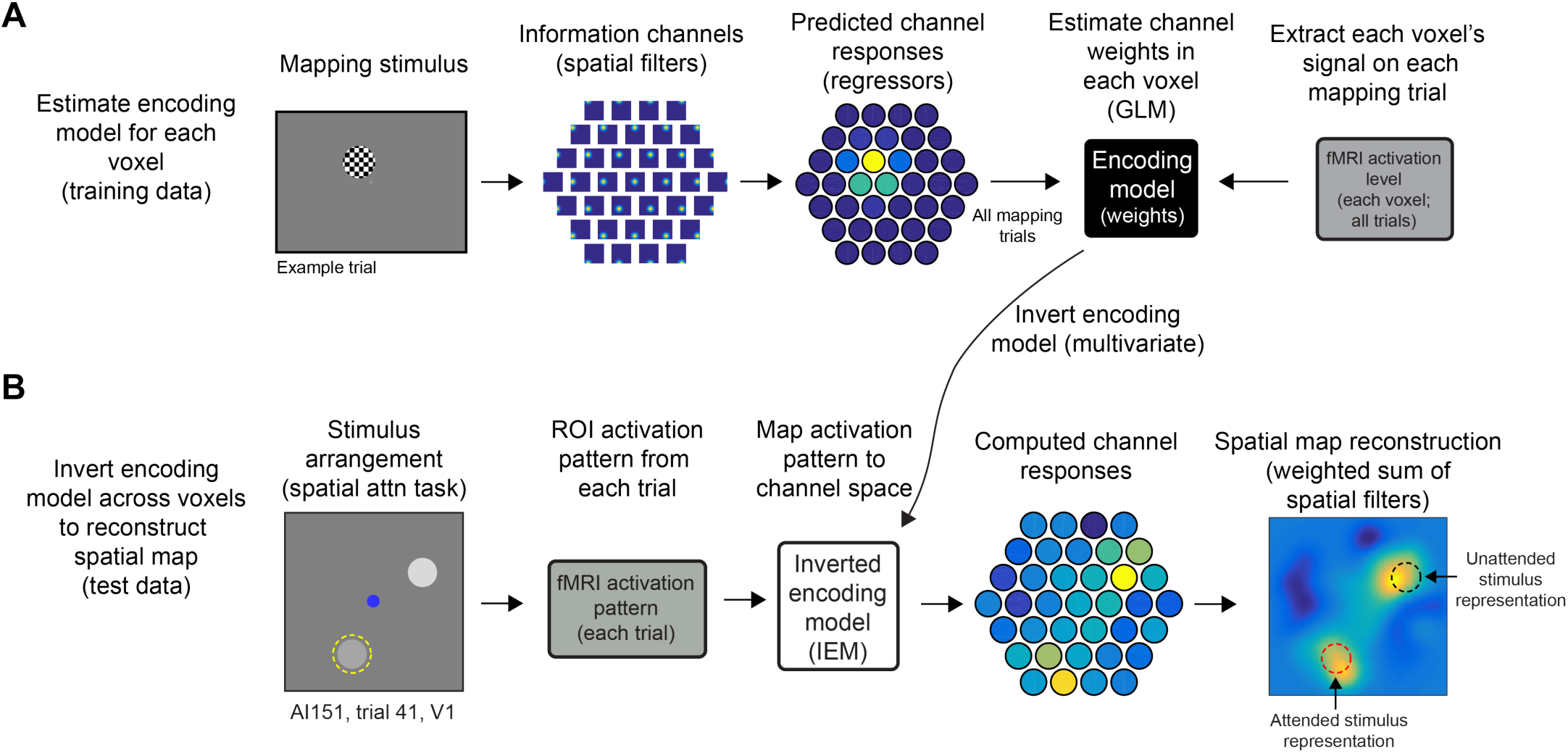
Measuring salience and relevant maps with inverted encoding model. **(A)** We estimated a spatial encoding model using data from a spatial mapping task in which participants viewed flickering checkerboard discs presented across a triangular grid of positions spanning a hexagonal region of the screen with spatial jitter (see (Sprague et al., 2016)). We modeled each voxel’s spatial sensitivity (i.e., receptive field or RF) as a weighted sum of smooth functions centered at each point in a triangular grid spanning the stimulus display visible inside the scanner. By knowing the stimulus on each trial (contrast mask) and the sensitivity profile of each modeled spatial ‘information channel’ (spatial filter), we could predict how each modeled information channel should respond to each stimulus. We then solve for the contribution of each channel to the measured signal in each voxel. This step amounts to solving the standard general linear model typical in fMRI analyses, and is univariate (each voxel can be estimated independently). **(B)** Then, we used all voxels within a region to compute an *inverted encoding model* (IEM). This step is multivariate; all voxels contribute to the IEM. This IEM allows us to transform activation patterns measured during the main spatial attention task (**Fig. 2A**) to activations of modeled information channels. Then, we compute a weighted sum of information channels based on their computed activation in the spatial attention task. These images of the visual field index the activation across the entire cortical region transformed into visual field coordinates. To quantify activation in the map, we extract the mean signal over the map area corresponding to the known stimulus positions on that trial (red dashed circle: attended stimulus position; black dashed circle: unattended stimulus position). For visualization of spatial maps averaged across trials, we rotated reconstructions as though stimuli were presented at positions indicated in cartoons (e.g., **Fig. 2B-C**). See (Sprague et al., 2016) for detailed methods on image alignment and coregistration.

This triangular grid of filters forms the set of information channels for our analysis. Each mapping task stimulus is converted from a contrast mask (1’s for each pixel subtended by the stimulus, 0’s elsewhere) to a set of filter activation levels by taking the dot product of the vectorized stimulus mask and the sensitivity profile of each filter. This results in each mapping stimulus being described by 37 filter activation levels rather than 1,024 × 768 = 786,432 pixel values. Once all filter activation levels are estimated, we normalize so that the maximum filter activation is 1.

Following previous reports (Brouwer and Heeger, 2009; Sprague and Serences, 2013), we model the response in each voxel as a weighted sum of filter responses (which can loosely be considered as hypothetical discrete neural populations, each with spatial RFs centered at the corresponding filter position).

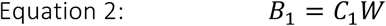

Where *B*_*1*_ (*n* trials × *m* voxels) is the observed BOLD activation level of each voxel during the spatial mapping task (averaged over two TRs, 6.00-8.00 s after mapping stimulus onset), *C*_*1*_ (*n* trials × *k* channels) is the modeled response of each spatial filter, or information channel, on each non-target trial of the mapping task (normalized from 0 to 1), and *W* is a weight matrix (*k* channels × *m* voxels) quantifying the contribution of each information channel to each voxel. Because we have more stimulus positions than modeled information channels, we can solve for *W* using ordinary least-squares linear regression:

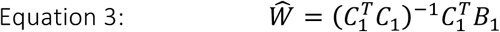

This step is univariate and can be computed for each voxel in a region independently. Next, we used all estimated voxel encoding models within a ROI (Ŵ) and a novel pattern of activation from the WM task (each TR from each trial, in turn) to compute an estimate of the activation of each channel (Ĉ_2_, *n* trials × *k* channels) which gave rise to that observed activation pattern across all voxels within that ROI (*B*_*2*_*, n* trials × *m* voxels):

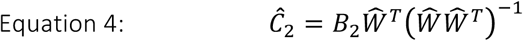

Once channel activation patterns are computed (Equation 4), we compute spatial reconstructions by weighting each filter’s spatial profile by the corresponding channel’s reconstructed activation level and summing all weighted filters together. This step aids in visualization, quantification, and coregistration of trials across stimulus positions, but does not confer additional information.

We used 4 mapping task runs to estimate the encoding model for each voxel, then inverted that encoding model to reconstruct visual field images during all main spatial attention task runs.

Because stimulus positions were unique on each trial of the selective attention task (**Fig. 2C**), direct comparison of image reconstructions on each trial is not possible without coregistration of reconstructions so that stimuli appeared at common positions across trials. To accomplish this, we adjusted the center position of the spatial filters on each trial such that we could rotate the resulting reconstruction. For **Figure 4**, we rotated each trial such that one target (the non-attended stimulus, **Fig. 2B**) was centered at *x* = 3.5° and *y* = 0° and the other stimulus was in the upper visual hemifield, which required flipping 1/2 of reconstructions across the horizontal meridian.

**Figure 4:**
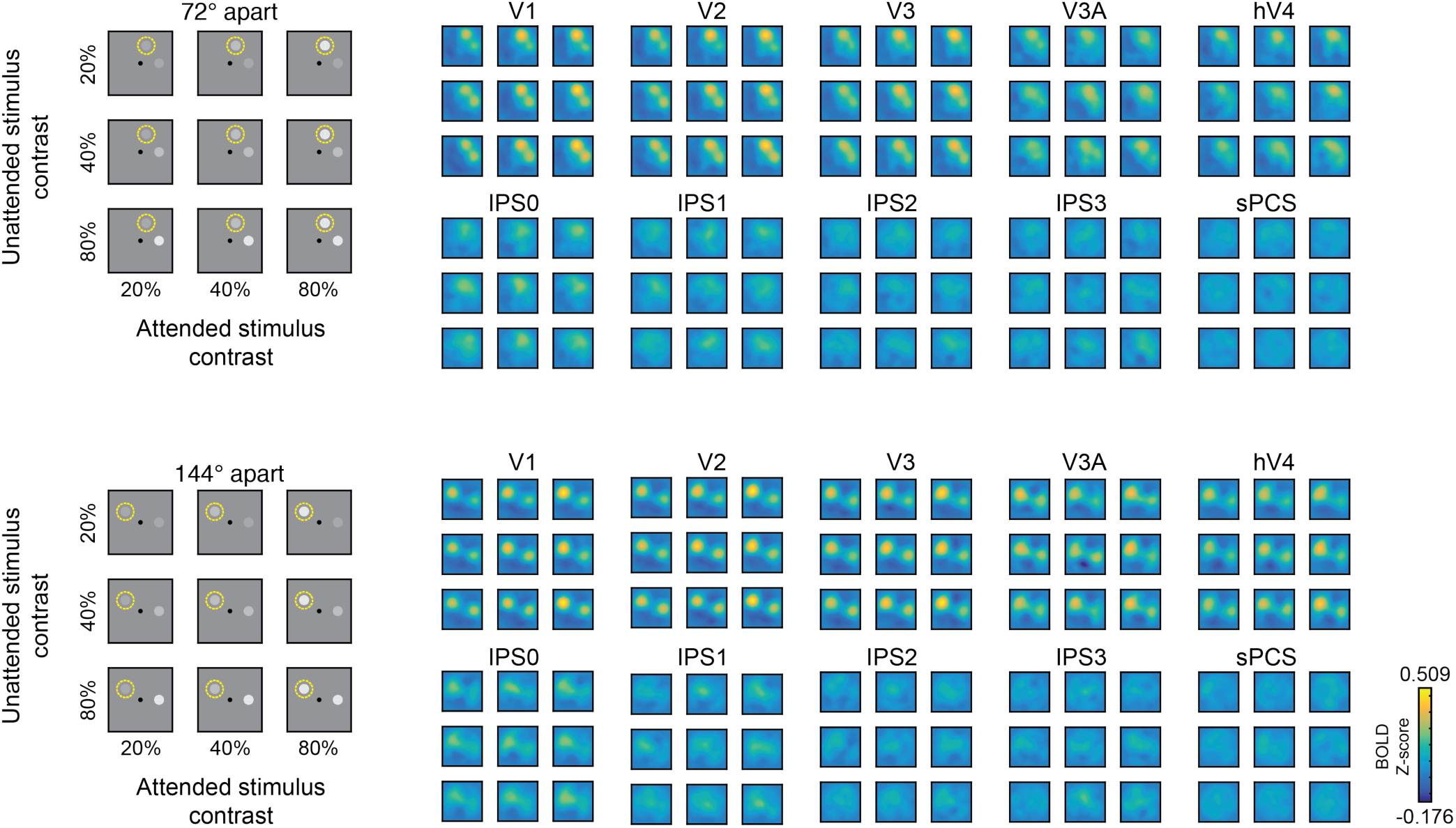
Reconstructed spatial maps index stimulus salience and relevance. Image reconstructions computed for each stimulus separation condition (grouped rows) and each stimulus contrast pair (entries within each 3x3 group of reconstructions). Within each group of reconstructions, the diagonal (top left to bottom right) are trials with matched contrasts, and are useful for visualizing the qualitative effect of behavioral relevance (spatial attention) on map profiles. Even in V1, locations near attended stimuli are represented more strongly than those near unattended stimuli. Going top-to-bottom (unattended stimulus) and left-to-right (attended stimulus) within each group of reconstructions steps through increasing stimulus contrast levels, and can be used to infer the sensitivity of each visual field map to visual salience.

### Quantifying Stimulus Representations

To quantify the strength of stimulus representations within each reconstruction, we averaged the pixels within each reconstruction located within a 0.9° radius disc centered at each stimulus’ known position. This gives us a single value for each stimulus (attended & unattended) on each trial. We then sorted these measurements, which we call “map activation” values as they reflect linear transformations of BOLD activation levels, based on the contrast of the attended and the unattended stimuli (**Figs. 5-6**).

**Figure 5:**
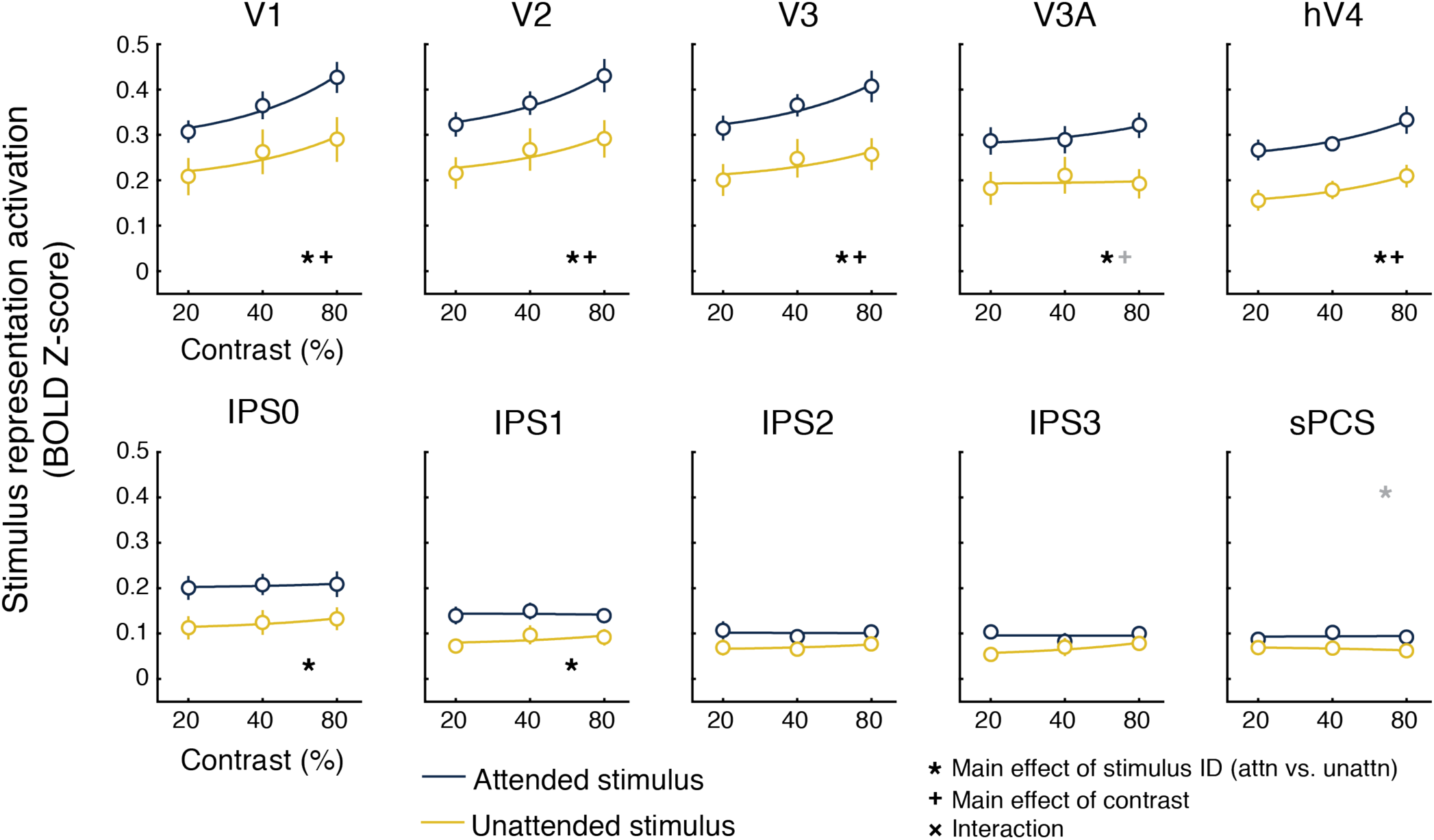
Quantifying sensitivity to visual salience and behavioral relevance across cortex. For each stimulus, we averaged the reconstruction activation on all trials where each stimulus appeared at a given contrast (see also Fig. 6). In V1, V2, V3, and hV4, reconstructed map activation increased with stimulus salience, regardless of whether the stimulus was attended. In all ROIs except for IPS2, IPS3, and sPCS, reconstructed map activation also reflected behavioral relevance, such that the attended location was more active than the unattended location. Error bars SEM across participants (*n* = 8). Lines indicate best-fit linear contrast response function, and all data is plotted on a log scale. All *p*-values available in **Extended data table 5-1**. Black symbols indicate significant main effects or interaction (FDR-corrected across all ROIs), and gray symbols indicate trend (*p* ≤ 0.05, uncorrected).

### Statistical Analysis

For all statistical tests, we used parametric tests (repeated measures ANOVAs and *T*-tests, where appropriate), followed up by 1,000 iterations of a randomized version of the test to derive an empirical null statistic distribution, given our data, from which we compute *p*-values reported throughout the text. If our limited sample satisfies the assumptions of these parametric tests, the *p*-values derived from the empirical null statistic distribution should closely approximate the derived value given assumptions. Especially because of our relatively small sample size, we prefer to rely on the empirical null for recovering *p*-values.

For behavioral analyses (**Fig. 2**), we computed a 2-way repeated-measures analysis of variance (ANOVA) with attended stimulus contrast and unattended stimulus contrast as factors for each of behavioral accuracy and response time. As a first neural analysis, to determine whether it was possible to collapse over sets of trials in which the irrelevant stimulus contrast varied (see **Fig. 6**), we computed a 2-way repeated-measures ANOVA with attended stimulus contrast and unattended stimulus contrast as factors for each of the stimulus representation activation values for each ROI. We were primarily interested in whether there were any interactions between attended and unattended stimulus contrast, which would have precluded us from collapsing over non-sorted stimulus contrasts. For completeness, *p*-values from our shuffling procedure for both main effects and the interaction for each ROI are presented in the Extended Data Tables 5-1 and 6-1. For a subsequent neural analysis testing the effects of salience and relevance on map activation across ROIs, we first conducted a 3-way ANOVA with factors of stimulus contrast, stimulus identity (attended vs. unattended), and ROI to identify whether there was a difference in attention-related changes in stimulus reconstructions across ROIs (as indicated by interactions between ROI and any other factor). Then, we performed a follow-up analysis on each ROI by computing a 2-way ANOVA with factors of stimulus contrast and stimulus identity (attended vs unattended) for each ROI.

**Figure 6:**
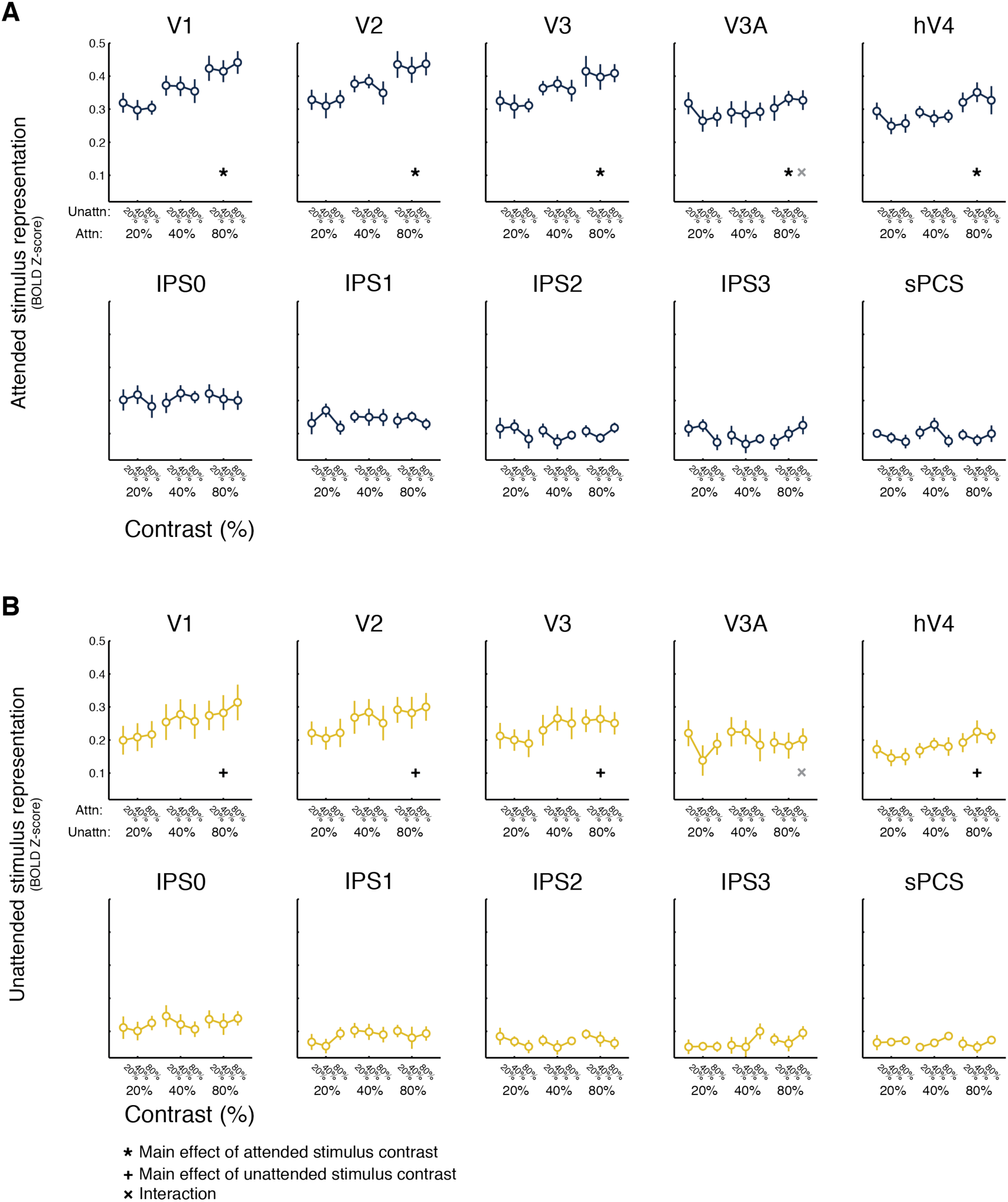
Attended stimulus representation does not depend on distractor stimulus contrast. **(A)** Reconstructed map activation for attended stimulus location across all salience combinations. Within each plot, linked points correspond to the contrast of the attended stimulus, and the individual linked points indicate the attended stimulus activation at each unattended stimulus contrast level (20%, 40%, 80%, left-to-right). Error bars SEM across participants. We observed no significant interactions (2-way permuted repeated-measures ANOVA within each ROI, factors of attended and unattended stimulus contrast, *p* ≥ 0.041, minimum *p*-value V3A), so we collapse across unattended stimulus salience within each attended stimulus salience level (**Fig. 5**). **(B)** Same, but for unattended stimulus activation, as sorted by attended stimulus contrast within each unattended stimulus contrast level. Again, we did not observe any significant interactions (*p* ≥ 0.026, minimum *p*-value V3A) so these are collapsed (**Fig. 5**). Error bars SEM. All *p*-values available in **Extended Data Table 6-1**. Black symbols indic ate significant main effects or interactions, FDR-corrected for multiple comparisons across ROIs. Gray symbols indicate trends, defined as *p* ≤ 0.05, uncorrected.

On each iteration of our shuffling procedure, we shuffled the data values for each participant individually and recomputed the test statistic of interest. To derive p-values, we computed the percentage of shuffling iterations in which the ‘null’ test statistic was greater than or equal to the measured test statistic (with intact, unshuffled labels). The random number generator was seeded with a single value for all analyses (derived by asking a colleague for a random string of numbers over instant messenger). For ROI analyses, trials were shuffled identically for each ROI. When appropriate (**Figs. 5-6**), we controlled for multiple comparisons using the false discovery rate (Benjamini and Yekutieli, 2001) across all comparisons within an analysis. All error bars reflect standard error of the mean, unless indicated otherwise.

### Data and Code Availability

All data and stimulus presentation and data analysis code necessary to produce figures supporting findings reported here [will be made upon acceptance or request from editor/reviewer] freely accessible online in an Open Science Framework repository (osf.io/XXXX), and code [will also be] maintained on github at author TCS’s profile (github.com/tommysprague/XXXX).

### Results

We measured blood oxygenation level dependent (BOLD) activation patterns from each independently-identified retinotopic region using functional magnetic resonance imaging (fMRI) while participants covertly attended one of two visual stimuli (two patches of randomly oriented dark and light lines), each presented at one of three luminance contrast levels (20%, 40%, and 80%; **Fig. 2A-B**), to identify a brief target stimulus (coherent lines that formed a spiral) at the attended location. We maintained behavioral performance at a constant accuracy level of ~80% (**Fig. 2D;** 2-way permuted repeated-measures ANOVA, *p*-values for main effect of attended, unattended contrast and interaction: 0.359, 0.096, and 0.853, respectively), so that any activation changes we observed did not reflect differences in task difficulty or engagement across experimental conditions. Additionally, response time did not vary with attended or distractor stimulus contrast, or their interaction (**Fig. 2E**; *p* = 0.926, 0.143, and 0.705, respectively), and the threshold coherence did not vary with the attended stimulus contrast (**Fig. 2F**; 1-way permuted repeated-measures ANOVA, main effect of attended contrast, *p* = 0.153).

We used a multivariate fMRI image reconstruction technique (an *inverted encoding model* or *IEM*) to visualize spatial maps of the visual scene using activation patterns from several cortical regions in visual, parietal and fontal cortex (Sprague and Serences, 2013). First, we estimated the spatial sensitivity profile of each voxel within a region of interest (ROI) using data measured from a separate set of ‘mapping’ scans. Then, we used the resulting sensitivity profiles across all voxels to reconstruct a map of retinotopic space in visual field coordinates from single-trial activation patterns measured during the covert visual attention task (**Fig. 3**). The spatial profile of activation within these maps can be used to infer whether a given ROI is sensitive to visual salience (i.e., does the spatial profile scale with contrast?, **Fig. 1B**) and whether it is sensitive to behavioral relevance (i.e., does the spatial profile scale with attention?, **Fig. 1D**). In principle, both visual salience and behavioral relevance could independently alter the landscape of responses in each ROI.

We found that image reconstructions systematically tracked the locations of stimuli in the visual field (**Fig. 4**). Qualitatively, the reconstructions from primary visual cortex (V1) reflected both stimulus salience and behavioral relevance: the reconstructed map activations at stimulus locations scaled both with increasing contrast and with the behavioral relevance of each stimulus. This pattern can be seen in **Fig. 4:** along the diagonal, where visual salience is equal between the two stimuli, map locations near the attended stimulus are more strongly active than locations near the unattended stimulus. However, in posterior parietal cortex (IPS0), only locations near the attended location were substantially active, with little activation associated with the irrelevant item’s location. Additionally, map activation in posterior parietal cortex did not scale with stimulus contrast. Importantly, even when the unattended stimulus was much more salient than the attended stimulus, only the attended stimulus location was strongly active. This demonstrates that behavioral relevance dominates activation profiles in this area. Overall, occipital retinotopic ROIs (V1-hV4, V3A) show a qualitative pattern similar to that observed in V1, and parietal and frontal ROIs show a pattern similar to that in IPS0.

To quantify these effects, we extracted the mean activation level in each reconstructed map at the known position of each stimulus and then evaluated the main effect of visual salience, the main effect of behavioral relevance, and their interaction using a repeated-measures 2-way ANOVA (with *p*-values computed using a randomization test and corrected for multiple comparisons via the false discovery rate, FDR, see **Materials & Methods: Statistical analyses**). If visual salience and behavioral relevance independently contribute to representations of attentional priority, we would expect to find a main effect of salience (contrast) and/or relevance (attention) on map activation in any given ROI, but no interactions between the two.

Reconstructed map activation increased significantly with visual salience in V1-hV4 (**Fig. 5**; *p* ≤ 0.003; see also **Fig. 6**), with no evidence for sensitivity to salience in parietal or frontal regions. On the other hand, map activation increased significantly with behavioral relevance not only in V1-hV4, but also in V3A and IPS0-1 (*p* ≤ 0.011), with a trend observed in sPCS (*p* = 0.038, trend defined at α = 0.05 without correction for multiple comparisons). There was no interaction between salience and relevance in any visual area that we evaluated (*p* ≥ 0.062, minimum *p*-value for V1). Additionally, a 3-way repeated-measures ANOVA with factors for salience, relevance, and ROI established that the influence of salience and relevance on map activation significantly varied across ROIs (interaction between salience and ROI: *p* < 0.001, interaction between relevance and ROI: *p* < 0.001; all *p*-values for 2- and 3-way ANOVAs available in Extended Data **Table 5-1**).

We also tested whether the map activation near one stimulus location depended on the contrast of the other stimulus. A 2-way repeated-measures ANOVA within each ROI on attended and unattended stimulus map activation, each with factors of attended and unattended contrast, yielded no significant interactions (*p* ≥ 0.124, minimum p-value sPCS; though trends emerged in V3A, defined as α = 0.05 without correction for multiple comparisons; **Fig. 6**; all *p*-values available in Extended Data **Table 6-1**). Thus, the unattended stimulus contrast did not impact reconstructed map activation near the attended stimulus, and vice versa.

## Discussion

We tested the hypothesis that attentional priority is guided by dissociable neural representations of the visual salience and the behavioral relevance of different spatial locations in the visual scene. While previous work has demonstrated sensitivity to bottom-up stimulus salience and/or top-down behavioral relevance in visual (Bogler et al., 2011, 2013; Burrows and Moore, 2009; Pestilli et al., 2011; Poltoratski et al., 2017; Somers et al., 1999; Zhang et al., 2012), parietal (Balan and Gottlieb, 2006; Bisley and Goldberg, 2003; Constantinidis and Steinmetz, 2005; Gottlieb et al., 1998; Serences and Yantis, 2007) and frontal (Bichot and Schall, 1999; Serences and Yantis, 2007; Thompson et al., 1997) cortex, it remains unexplored how each of these factors interact with one another when parametrically manipulated, as well as how large-scale priority maps subtended by entire visual retinotopic regions support representations of salience, relevance, and their interaction. Here, we identified representations of visual salience within reconstructed attentional priority maps measured from early visual regions, where map activation scaled with image-computable visual salience. Additionally, we identified representations of behavioral relevance across visual and posterior parietal cortex, with some regions, such as IPS0, exhibiting activation profiles which only index the location of a relevant object, even when an irrelevant but salient object was simultaneously present in the display.

These results provide evidence for a distributed representation of attentional priority, with each retinotopic region supporting a combination of independent representations of the visual salience and behavioral relevance of scene elements (Fecteau and Munoz, 2006; Itti and Koch, 2001; Serences and Yantis, 2006). Models of attentional priority have implemented layers corresponding to image-computable visual salience and task-driven behavioral relevance, but large-scale spatial maps of such features have only been documented in isolation (i.e., manipulations of relevance given equal image-computable salience (Jerde et al., 2012; Serences and Yantis, 2007; Sprague and Serences, 2013), or of salience given equal behavioral relevance (Bogler et al., 2011, 2013; White et al., 2017; Zhang et al., 2012)) or at a local physiological scale (i.e., measurements of single neurons in isolation during manipulation of item salience and/or relevance (Bichot and Schall, 1999; Bisley and Goldberg, 2003, 2010; Burrows and Moore, 2009; Gottlieb et al., 1998; Ipata et al., 2006; Mazer and Gallant, 2003; Thompson et al., 1997; White et al., 2017)). Here, we independently manipulated the salience and relevance of multiple scene elements and reconstructed large-scale spatial maps using all voxels within a retinotopic region. Because spatial attention has nuanced and multifaceted effects on voxels tuned to locations across the visual field (de Haas et al., 2014; Kay et al., 2015; Klein et al., 2014; Sheremata and Silver, 2015; Sprague and Serences, 2013; Vo et al., 2017), it was necessary to apply the population-level IEM analysis technique to measure the joint effect of all task-related modulations on spatial maps supported by entire regions. This allowed us to demonstrate that salience and relevance each independently contribute to the representation of attentional priority, and that maps become more sensitive to relevance and less sensitive to salience across the visual processing hierarchy (**Fig. 5**).

In our stimulus setup, salience could only be defined by the luminance contrast of each stimulus. Other studies have defined stimulus salience based on sudden stimulus onsets, or based on distinct stimulus features among a field of distractors (‘singletons’). In these previous studies, neural activity in macaque LIP and FEF exhibited properties consistent with a salience map: neurons respond to abrupt-onset stimuli, but not to stable features of the environment (Balan and Gottlieb, 2006; Gottlieb et al., 1998; Kusunoki et al., 2000), and they respond strongly to singleton items among uniform distractors (Bichot and Schall, 1999; Thompson et al., 1997). While we did not see responses in parietal cortex consistent with an image-computable salience map (**Fig. 4-5**), it remains possible that other stimulus manipulations could result in salience representations in parietal and frontal cortex. A recent fMRI study presented participants with orientation or motion singletons and found responses consistent with salience maps in visual cortex, similar to our observations here, although they did not report any parietal cortex data (Poltoratski et al., 2017).

Indeed, results from other studies suggest that the type of salience manipulation may have a substantial impact on modulations measured in humans with fMRI. In a study in which local contrast dimming among patterned patches was used to define a salient region, activation in early visual field maps reflected the decrease in contrast (but increase in salience by virtue of the high-contrast background) with a decrease in activation, rather than the increase in activation expected if these regions detect salience (Betz et al., 2013). This conflicts with another study in which briefly-presented and masked orientation texture patches induced salience-related responses in early visual cortex (Zhang et al., 2012; but see Bogler et al., 2013). Finally, in our study, we only observed salience-related activation profiles in occipital regions of visual cortex (**Figs. 4-5**; V1-hV4). Future studies should compare how different types of bottom-up salience manipulations alter activation profiles of putative priority maps across visual, parietal, and frontal cortex.

Additionally, many other studies examining the interaction between behavioral relevance and visual salience use visual search tasks, in which the locus of spatial attention must explore the visual scene on each trial to identify a target stimulus. For example, an animal or human may be required to report the orientation of a bar presented within a green square among a field of bars, each surrounded by green circles. As a salient distractor, one circle might be red. In such a task, the relevant location cannot be known ahead of time by the subject. We designed our study to parametrically manipulate the spatial location of covert attention via an endogenous cue (Bisley and Goldberg, 2003; Carrasco, 2011), as well as the location of a distracting stimulus of varying levels of salience. Accordingly, we could sort trials based on the location where attention was instructed, which allowed us to quantify changes in map activation levels according to behavioral relevance. When attention is free to roam during a visual search task, all locations may be sampled during a trial, which would be impossible to pull apart using hemodynamic measures. However, recent application of IEM analyses to visual covert attention tasks may enable high-temporal-resolution studies into the wandering spotlight of attention during such visual search tasks (Foster et al., 2017; Garcia et al., 2013; Samaha et al., 2016).

The relative balance of stimulus salience and behavioral relevance within a region may shift under additional task conditions, such as manipulations of cue validity or reward magnitude (Chelazzi et al., 2014; Kahnt et al., 2014; Klyszejko et al., 2014), or when participants are cued to attend to different feature values rather than spatial positions. We speculate that when behavioral goals are ill-defined, maps that are more sensitive to image-computable salience will primarily determine attentional priority and guide behavior (Egeth and Yantis, 1997). In contrast, when an observer is highly focused on a specific behavioral goal, maps with stronger representations of behavioral relevance will dominate the computation of attentional priority.

## Author Contributions

TCS, SI, VAV, and JTS conceived and designed the research. TCS, SI, and VAV acquired data, TCS analyzed data and wrote the first draft of the manuscript, SI, VAV and JTS provided comments & critical edits to the manuscript. JTS supervised all aspects of the research.

## Acknowledgements

This work was supported by NEI R01-EY025872 and a James S McDonnell Foundation Scholar Award to JTS, a HHMI Graduate Student Fellowship to SI, and a NSF Graduate Research Fellowship to VAV. We thank Rosanne Rademaker, Edward Ester, Miranda Scolari, Bradley Postle, Clayton Curtis, Kartik Sreenivasan, and Kevin DeSimone for helpful discussions.

---

**Extended Data Table 5-1:** P-VALUES FOR ALL TESTS IN Figure 5. To assess whether each ROI was sensitive to visual salience and behavioral relevance (and their interaction), we performed a 2-way, repeated-measures ANOVA with factors of visual salience (20%, 40%, 80% contrast) and behavioral relevance (attended or unattended). To generate *p*-values, we compared the *F*-score derived for each main effect and their interaction to those derived from a shuffling procedure in which we shuffled the data labels within each participant independently 1,000 times. Because we ran 1,000 iterations of this shuffling procedure, the minimum accurate quantifiable p-value was 0.001. Bold p-values indicate significant effects after correcting for multiple comparisons (false discovery rate, *q* = 0.05 across all comparisons, threshold *p* ≤ 0.011). Italics p-values indicate trends, defined as *p* ≤ 0.05, no correction for multiple comparisons. Int: Interactions.

|  | V1 | V2 | V3 | V3A | hV4 | IPSO | IPS1 | IPS2 | IPS3 | sPCS |
| --- | --- | --- | --- | --- | --- | --- | --- | --- | --- | --- |
| Saliency | **0.003** | **<0.001** | **<0.001** | *0.022* | **<0.001** | 0.223 | 0.394 | 0.371 | 0.653 | 0.717 |
| Relevance | **<0.001** | **0.002** | **<0.001** | **0.003** | **0.005** | **0.001** | **0.011** | 0.071 | 0.112 | *0.038* |
| Interaction | 0.062 | 0.183 | 0.117 | 0.166 | 0.291 | 0.755 | 0.527 | 0.922 | 0.204 | 0.644 |
| **Three-way ANOVA** | **Main effects** | Saliency | Relevance | ROI | **Int.** | Saliency × Relevance | Saliency × ROI | Relevance × ROI | Saliency × Relevance × ROI |  |
|  |  | **<0.001** | **0.001** | **<0.001** |  | 0.348 | **<0.001** | **<0.001** | 0.166 |  |

**Extended Data Table 6-1:**
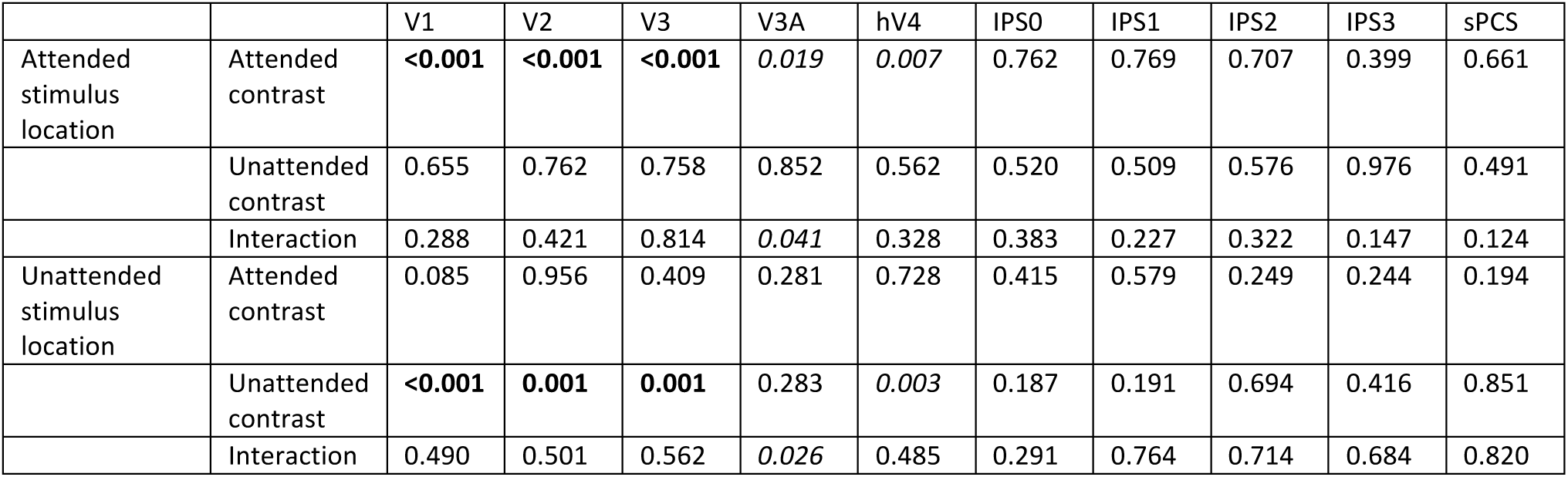
P-VALUES FOR ALL TESTS REPORTED IN Fig. 6. To test whether the map activation at one stimulus location depended on the contrast of the stimulus presented at the other location, we conducted a set of 2-way repeated-measures ANOVAs. First, we tested whether the attended stimulus location map activation varied as a function of attended stimulus contrast (20%, 40%, 80%), unattended stimulus contrast (20%, 40%, 80%), or their interaction (top 3 rows). Then, we tested whether the unattended stimulus location map activation varied as a function of attended stimulus contrast, unattended stimulus contrast, and their interaction (bottom 3 rows). All p-values were computed against null tests in which we shuffled condition labels within each participant independently 1,000 times and compared the intact *F*-values against those derived from the shuffled distribution. Because we ran 1,000 iterations of this shuffling procedure, the minimum accurate quantifiable p-value was 0.001. We corrected for multiple comparisons across all tests via FDR (bold). Italics indicates trends, defined as *p* ≤ 0.05, uncorrected.

|  |  | V1 | V2 | V3 | V3A | hV4 | IPSO | IPS1 | IPS2 | IPS3 | sPCS |
| --- | --- | --- | --- | --- | --- | --- | --- | --- | --- | --- | --- |
| Attended stimulus location | Attended contrast | **<0.001** | **<0.001** | **<0.001** | *0.019* | *0.007* | 0.762 | 0.769 | 0.707 | 0.399 | 0.661 |
|  | Unattended contrast | 0.655 | 0.762 | 0.758 | 0.852 | 0.562 | 0.520 | 0.509 | 0.576 | 0.976 | 0.491 |
|  | Interaction | 0.288 | 0.421 | 0.814 | *0.041* | 0.328 | 0.383 | 0.227 | 0.322 | 0.147 | 0.124 |
| Unattended stimulus location | Attended contrast | 0.085 | 0.956 | 0.409 | 0.281 | 0.728 | 0.415 | 0.579 | 0.249 | 0.244 | 0.194 |
|  | Unattended contrast | **<0.001** | **0.001** | **0.001** | 0.283 | *0.003* | 0.187 | 0.191 | 0.694 | 0.416 | 0.851 |
|  | Interaction | 0.490 | 0.501 | 0.562 | *0.026* | 0.485 | 0.291 | 0.764 | 0.714 | 0.684 | 0.820 |

## References

Balan, P.F., and Gottlieb, J. (2006). Integration of exogenous input into a dynamic salience map revealed by perturbing attention. J. Neurosci. 26, 9239–9249.

Benjamini, Y., and Yekutieli, D. (2001). The control of the false discovery rate in multiple testing under dependency. Ann. Stat. 29, 1165–1188.

Bertleff, S., Fink, G.R., and Weidner, R. (2016). The Role of TopDown Focused Spatial Attention in Preattentive Salience Coding and Salience-based Attentional Capture. J. Cogn. Neurosci. 28, 1152–1165.

Betz, T., Wilming, N., Bogler, C., Haynes, J.-D., and König, P. (2013). Dissociation between saliency signals and activity in early visual cortex. J. Vis. 13, 6–6.

Bichot, N.P., and Schall, J.D. (1999). Effects of similarity and history on neural mechanisms of visual selection. Nat Neurosci 2, 549–554.

Bisley, J.W., and Goldberg, M.E. (2003). Neuronal activity in the lateral intraparietal area and spatial attention. Science (80.). 299, 81–86.

Bisley, J.W., and Goldberg, M.E. (2010). Attention, intention, and priority in the parietal lobe. Annu. Rev. Neurosci. 33, 1–21.

Bogler, C., Bode, S., and Haynes, J.-D. (2011). Decoding Successive Computational Stages of Saliency Processing. Curr. Biol. 21, 1667–1671.

Bogler, C., Bode, S., and Haynes, J.-D. (2013). Orientation pop-out processing in human visual cortex. Neuroimage 81, 73–80.

Brainard, D.H. (1997). The Psychophysics Toolbox. Spat. Vis. 10, 433–436.

Brouwer, G., and Heeger, D. (2009). Decoding and Reconstructing Color from Responses in Human Visual Cortex. J. Neurosci. 29, 13992–14003.

Burrows, B.E., and Moore, T. (2009). Influence and limitations of popout in the selection of salient visual stimuli by area V4 neurons. J. Neurosci. 29, 15169–15177.

Carrasco, M. (2011). Visual attention: the past 25 years. Vision Res. 51, 1484–1525.

Chelazzi, L., Exštočinovà, J., Calletti, R., Lo Gerfo, E., Sani, I., Della Libera, C., and Santandrea, E. (2014). Altering spatial priority maps via reward-based learning. J. Neurosci. 34, 8594–8604.

Constantinidis, C., and Steinmetz, M.A. (2005). Posterior parietal cortex automatically encodes the location of salient stimuli. J. Neurosci. 25, 233–238.

Egeth, H.E., and Yantis, S. (1997). Visual attention: control, representation, and time course. Annu. Rev. Psychol. 48, 269–297.

Engel, S.A., Rumelhart, D.E., Wandell, B.A., Lee, A.T., Glover, G.H., Chichilnisky, E.-J., and Shadlen, M.N. (1994). fMRI of human visual cortex. Nature 369, 525.

Ester, E.F., Sprague, T.C., and Serences, J.T. (2015). Parietal and Frontal Cortex Encode Stimulus-Specific Mnemonic Representations during Visual Working Memory. Neuron 87, 893–905.

Fecteau, J.H., and Munoz, D.P. (2006). Salience, relevance, and firing: a priority map for target selection. Trends Cogn. Sci. 10, 382–390.

Folk, C.L., Remington, R.W., and Johnston, J.C. (1992). Involuntary covert orienting is contingent on attentional control settings. J. Exp. Psychol. Hum. Percept. Perform. 18, 1030–1044.

Folk, C.L., Leber, A.B., and Egeth, H.E. (2002). Made you blink! Contingent attentional capture produces a spatial blink. Percept. Psychophys. 64, 741–753.

Foster, J.J., Sutterer, D.W., Serences, J.T., Vogel, E.K., and Awh, E. (2017). Alpha-Band Oscillations Enable Spatially and Temporally Resolved Tracking of Covert Spatial Attention. Psychol. Sci. 28, 929–941.

Freeman, J., Brouwer, G.J., Heeger, D.J., and Merriam, E.P. (2011). Orientation decoding depends on maps, not columns. J. Neurosci. 31, 4792–4804.

Freeman, J., Heeger, D.J., and Merriam, E.P. (2013). Coarse-scale biases for spirals and orientation in human visual cortex. J. Neurosci. 33, 19695–19703.

Garcia, J., Srinivasan, R., and Serences, J. (2013). Near-Real-Time Feature-Selective Modulations in Human Cortex. Curr. Biol. 23, 515–522.

Gottlieb, J.P., Kusunoki, M., and Goldberg, M.E. (1998). The representation of visual salience in monkey parietal cortex. Nature 391, 481–484.

de Haas, B., Schwarzkopf, D.S., Anderson, E.J., and Rees, G. (2014). Perceptual load affects spatial tuning of neuronal populations in human early visual cortex. Curr. Biol. 24, R66–R67.

Hagler, D.J., and Sereno, M.I. (2006). Spatial maps in frontal and prefrontal cortex. Neuroimage 29, 567–577.

Ipata, A.E., Gee, A.L., Gottlieb, J., Bisley, J.W., and Goldberg, M.E. (2006). LIP responses to a popout stimulus are reduced if it is overtly ignored. Nat. Neurosci. 9, 1071–1076.

Itti, L., and Koch, C. (2001). Computational modelling of visual attention. Nat Rev Neurosci 2, 194–203.

Jerde, T.A., Merriam, E.P., Riggall, A.C., Hedges, J.H., and Curtis, C.E. (2012). Prioritized Maps of Space in Human Frontoparietal Cortex. J. Neurosci. 32, 17382–17390.

Kahnt, T., Park, S.Q., Haynes, J.-D., and Tobler, P.N. (2014). Disentangling neural representations of value and salience in the human brain. Proc. Natl. Acad. Sci. 111, 5000–5005.

Kay, K.N., Weiner, K.S., and Grill-Spector, K. (2015). Attention reduces spatial uncertainty in human ventral temporal cortex. Curr. Biol.

Klein, B.P., Harvey, B.M., and Dumoulin, S.O. (2014). Attraction of position preference by spatial attention throughout human visual cortex. Neuron 84, 227–237.

Klyszejko, Z., Rahmati, M., and Curtis, C.E. (2014). Attentional priority determines working memory precision. Vision Res. 105, 70–76.

Koch, C., and Ullman, S. (1985). Shifts in selective visual attention: towards the underlying neural circuitry. Hum. Neurobiol. 4, 219–227.

Kusunoki, M., Gottlieb, J., and Goldberg, M.E. (2000). The lateral intraparietal area as a salience map: the representation of abrupt onset, stimulus motion, and task relevance. Vision Res. 40, 1459–1468.

Mackey, W.E., Winawer, J., and Curtis, C.E. (2017). Visual field map clusters in human frontoparietal cortex. Life 6.

Masciocchi, C.M., Mihalas, S., Parkhurst, D., and Niebur, E. (2009). Everyone knows what is interesting: salient locations which should be fixated. J. Vis. 9, 25.122.

Mazer, J.A., and Gallant, J.L. (2003). Goal-related activity in V4 during free viewing visual search. Evidence for a ventral stream visual salience map. Neuron 40, 1241–1250.

Parkhurst, D., Law, K., and Niebur, E. (2002). Modeling the role of salience in the allocation of overt visual attention. Vision Res. 42, 107–123.

Pelli, D.G. (1997). The VideoToolbox software for visual psychophysics: transforming numbers into movies. Spat. Vis. 10, 437–442.

Pestilli, F., Carrasco, M., Heeger, D.J., and Gardner, J.L. (2011). Attentional enhancement via selection and pooling of early sensory responses in human visual cortex. Neuron 72, 832–846.

Poltoratski, S., Ling, S., McCormack, D., and Tong, F. (2017). Characterizing the effects of feature salience and top-down attention in the early visual system. J. Neurophysiol. 118, 564–573.

Rahmati, M., Saber, G.T., and Curtis, C.E. Population dynamics of early visual cortex during working memory. J. Cogn. Neurosci.

Saber, G.T., Pestilli, F., and Curtis, C.E. (2015). Saccade planning evokes topographically specific activity in the dorsal and ventral streams. J. Neurosci. 35, 245–252.

Samaha, J., Sprague, T.C., and Postle, B.R. (2016). Decoding and Reconstructing the Focus of Spatial Attention from the Topography of Alpha-band Oscillations. J. Cogn. Neurosci. 1–8.

Serences, J.T., and Yantis, S. (2006). Selective visual attention and perceptual coherence. Trends Cogn. Sci. 10, 38–45.

Serences, J.T., and Yantis, S. (2007). Spatially selective representations of voluntary and stimulus-driven attentional priority in human occipital, parietal, and frontal cortex. Cereb. Cortex 17, 284–293.

Sheremata, S.L., and Silver, M.A. (2015). Hemisphere-Dependent Attentional Modulation of Human Parietal Visual Field Representations. J. Neurosci. 35, 508–517.

Silver, M.A., Ress, D., and Heeger, D.J. (2005). Topographic maps of visual spatial attention in human parietal cortex. J. Neurophysiol. 94, 1358–1371.

Somers, D.C., Dale, A.M., Seiffert, A.E., and Tootell, R.B. (1999). Functional MRI reveals spatially specific attentional modulation in human primary visual cortex. Proc. Natl. Acad. Sci. U. S. A. 96, 1663–1668.

Sprague, T.C., and Serences, J.T. (2013). Attention modulates spatial priority maps in the human occipital, parietal and frontal cortices. Nat. Neurosci. 16, 1879–1887.

Sprague, T.C., Ester, E.F., and Serences, J.T. (2014). Reconstructions of Information in Visual Spatial Working Memory Degrade with Memory Load. Curr. Biol.

Sprague, T.C., Saproo, S., and Serences, J.T. (2015). Visual attention mitigates information loss in small and large-scale neural codes. Trends Cogn. Sci. 19, 215–226.

Sprague, T.C., Ester, E.F., and Serences, J.T. (2016). Restoring Latent Visual Working Memory Representations in Human Cortex. Neuron 91, 694–707.

Srimal, R., and Curtis, C.E. (2008). Persistent neural activity during the maintenance of spatial position in working memory. Neuroimage 39, 455–468.

Stigchel, S. Van der, Belopolsky, A. V., Peters, J.C., Wijnen, J.G., Meeter, M., and Theeuwes, J. (2009). The limits of topdown control of visual attention. Acta Psychol. (Amst). 132, 201–212.

Swisher, J.D., Halko, M.A., Merabet, L.B., McMains, S.A., and Somers, D.C. (2007). Visual topography of human intraparietal sulcus. J. Neurosci. 27, 5326–5337.

Theeuwes, J. (1994). Endogenous and exogenous control of visual selection. Perception 23, 429–440.

Theeuwes, J. (2010). Top–down and bottom–up control of visual selection. Acta Psychol. (Amst). 135, 77–99.

Thompson, K.G., Bichot, N.P., and Schall, J.D. (1997). Dissociation of visual discrimination from saccade programming in macaque frontal eye field. J. Neurophysiol. 77, 1046–1050.

Usher, M., and Niebur, E. (1996). Modeling the Temporal Dynamics of IT Neurons in Visual Search: A Mechanism for Top-Down Selective Attention. J. Cogn. Neurosci. 8, 311–327.

Veale, R., Hafed, Z.M., and Yoshida, M. (2017). How is visual salience computed in the brain? Insights from behaviour, neurobiology and modelling. Philos. Trans. R. Soc. B Biol. Sci. 372, 20160113.

Vo, V.A., Sprague, T.C., and Serences, J.T. (2017). Spatial Tuning Shifts Increase the Discriminability and Fidelity of Population Codes in Visual Cortex. J. Neurosci. 37, 3386–3401.

White, B.J., Berg, D.J., Kan, J.Y., Marino, R.A., Itti, L., and Munoz, D.P. (2017). Superior colliculus neurons encode a visual saliency map during free viewing of natural dynamic video. Nat. Commun. 8, 14263.

Wolfe, J.M. (1994). Guided Search 2.0 A revised model of visual search. Psychon. Bull. Rev. 1, 202–238.

Yantis, S., and Johnson, D.N. (1990). Mechanisms of attentional priority. J. Exp. Psychol. Hum. Percept. Perform. 16, 812–825.

Zhang, X., Zhaoping, L., Zhou, T., and Fang, F. (2012). Neural activities in v1 create a bottomup saliency map. Neuron 73, 183–192.

